# Structural Basis of Differential Gene Expression at eQTLs Loci from High-Resolution Ensemble Models of 3D Single-Cell Chromatin Conformations

**DOI:** 10.1101/2024.06.13.598877

**Authors:** Lin Du, Hammad Farooq, Pourya Delafrouz, Jie Liang

## Abstract

**Motivation:** Techniques such as high-throughput chromosome conformation capture (Hi-C) have provided a wealth of information on the organization of the nucleus and the genome important for understanding gene expression regulation. Additionally, Genome-Wide Association Studies (GWASs) have uncovered thousands of loci related to complex traits. Expression quantitative trait loci (eQTL) studies have further linked the genetic variants to alteration in expression levels of associated target genes across individuals. However, the functional roles of many eQTLs located in non-coding regions are unclear. Current joint analyses of Hi-C and eQTLs data lack advanced computational tools, limiting what can be learned from these data.

**Result:** In this work, we developed a computational method for simultaneous analysis of Hi-C and eQTL data. Our method can identify a small set of non-random interactions from all Hi-C interactions. Using these non-random interactions, we reconstruct large ensemble (×10^5^) of high-resolution single-cell 3D chromatin conformations with thorough sampling, which accurately replicate Hi-C measurements. Our results revealed the presence of many-body interactions in chromatin conformation at single-cell level in eQTL locus, offering detailed view into how three-dimensional structures of chromatin form the physical foundation for gene regulation, including how genetic variants of eQTLs affect the expression level of their associated eGenes.

Furthermore, our method can deconvolve chromatin heterogeneity and investigate the spatial associations of eQTLs and eGenes at subpopulation level to reveal their regulatory impacts on gene expression. Together, ensemble modeling of thoroughly sampled single cell chromatin conformations from Hi-C, along with eQTL data, helps to decipher how chromatin 3D structures provide the physical basis for gene regulation, expression control, and aid in understanding of the overall structure-function relationships of genome organization.

Availability and implementation: It is available at https://github.com/uic-liang-lab/3DChromFolding-eQTL-Loci

## 1. INTRODUCTION

The spatial organization of genome plays important roles in gene expression and DNA replication (Fudenberg et al. 2011, Sexton and Cavalli 2015, Pope et al. 2014). Studies based on chromosome conformation capture techniques (Baù and Marti-Renom 2012, Simonis et al. 2006, Dostie et al. 2006) such as Hi-C (Lieberman-Aiden et al. 2009) contained in the 4D Nucleome Project (the 4D Nucleome Network et al. 2017, Reiff et al. 2022), have provided a wealth of information on the 3D organization of genomes of many tissues and cell types. At the same time, a large number of genetic variants statistically associated with gene expression across individuals have been identified for 49 different tissues and are now available from the Genotype-Tissue Expression (GTEx) Consortium (Lonsdale et al. 2013, The GTEx Consortium et al. 2015, Aguet et al. 2017).

Joint analysis of eQTLs data and chromatin interactions has already revealed important insights (Aguet et al. 2017, Duggal, Wang and Kingsford 2014, Lu et al. 2020, Schmiedel et al. 2018, Yu, Hu and Li 2019). eQTLs are found to be spatially close to their target genes (Duggal, Wang and Kingsford 2014), and eQTLs enriched in cis regulatory elements tend to be in close spatial proximity with their target gene promoters (Aguet et al. 2017). A recent study showed that genes whose expression are significantly associated with eQTLs are positively correlated with chromatin contact frequencies. Further, eQTLs and their target genes are found to be more likely to co-localize within the same Topologically Associating Domains (TADs) (Yu, Hu and Li 2019).

However, current joint analyses that integrate Hi-C and eQTLs data have limitations. While there is abundant of Hi-C data, 3D spatial configurations of chromatin loci and interactions in individual cells cannot be directly inferred due to the intrinsic 2D and population-averaged nature of Hi-C frequency heatmaps, Hi-C data do not provide direct knowledge on how 3D structures of the chromatin determine gene expression in individual cells. Furthermore, it is also unclear that among the numerous chromatin interactions identified by Hi-C studies, which ones reflect functional associations, and which ones are due to by-stander effect, resulting from random collision owing to volume confinement and other factors (Belmont 2014, Gürsoy et al. 2014). In addition, many-body interactions in condensates involving more than pairwise (e.g. pairs of eQTLs and their associated genes, eGene-eQTLs) interactions play important roles in genome organization and function (Hnisz et al. 2017), but they cannot be directly identified from 2D Hi-C heatmaps. Furthermore, while heatmaps of population Hi-C exhibit highly detailed patterns, the heterogeneities of the chromatin 3D structures in the underlying cell population are difficult to assess. While there are several approaches for modeling 3D chromatin from experimental Hi-C data, including consensus optimization [ShRec3D (Lesne et al. 2014), ChromSDE (Zhang et al. 2013) and Chrom3D (Paulsen et al. 2017)], ensemble optimization [(Baù and Marti-Renom 2012), and PGS (Tjong et al. 2016, Hua et al. 2018)], and block copolymer models (Di Pierro et al. 2016, Zhang and Wolynes 2017, Conte et al. 2020, Contessoto, Cheng and Onuchic 2022) (reviewed in (Liang and Perez-Rathke 2021)), adequately sampling diverse chromatin conformations remains challenging (Liang and Perez-Rathke 2021). As chromatin conformations in a cell population may fall into a set of distinct structural clusters (Gürsoy et al. 2017, Sun et al. 2021), where each cluster harbors conformations of similar topology, some clusters may have favorable 3D spatial arrangements of promoter, enhancer, gene, and other elements to facilitate gene expression, while others may not. However, such chromatin heterogeneity and analysis of cell subpopulations with relevant chromatin structural clusters have not been quantified in the eQTLs enriched loci.

In this study, we develop a new computational method for simultaneous analysis of both Hi-C and eQTLs data, overcoming difficulties listed above. Specifically, we construct a pipeline for identifying non-random Hi-C interactions around eQTLs/eGenes that are beyond polymer collision. We further generate large ensembles (5×10^4^) of 3D single-cell chromatin conformations models for the selected locus. We then compute subpopulations of single-cell chromatin configurations at the locus and quantified the overall chromatin heterogeneity. Our tools allow interrogations of the 3D chromatin structures of loci where eQTLs and/or eGenes reside to gain an understanding of how they are spatially associated, and how such spacial relationship may affect regulation of gene expression. Our tools also identify the participating genes, promoters, and other elements in the spatial context of the eQTLs. Furthermore, with 3D ensemble models of their spatial arrangement constructed, our results uncovered higher-order many-body interacting units important for gene regulation of an eQTLs-rich locus across lymphoblastoid cells (GM12878), human primary mammary epithelial cells (HMEC), and lung fibroblast cells (IMR-90). Our results demonstrate the benefits of integrating Hi-C and eQTL data through deep sampling of large ensemble models of single cell chromatin conformations, allowing us to explore critical issues related to the differences in genome 3D structure across tissues and the relationship between genome 3D structure and function.

## 2 MATERIALS AND METHODS

### 2.1 Dataset

In this study, we used bulk Hi-C data of lymphoblastoid cells (GM12878, 4DNES3JX38V5), human primary mammary epithelial cells (HMEC, 4DNESIE5R9HS) and lung fibroblast cells (IMR-90, 4DNES1ZEJNRU) from the 4D Nucleome Data Portal (the 4D Nucleome Network et al. 2017, Reiff et al. 2022). We also obtained eQTLs data and bulk tissue expression data for cells of EBV-transformed lymphocytes, breast mammary tissue, and lung tissues from the GTEx Portal (Data Source: GTEx Analysis Release V8) (Lonsdale et al. 2013). All data are based on the reference genome GRCh38.

### 2.2 Overview of the pipeline

The overall computational pipeline of our method is illustrated in Figure 1. Our model builds on the recent advancements in methods of deep sampling to generate random polymer 3D chromatin conformations (Gürsoy et al. 2014, Gürsoy et al. 2017, Perez-Rathke et al. 2020, Sun et al. 2021, Perez-Rathke et al. 2019). We first use a sequential Monte Carlo approach (Gürsoy et al. 2014, Gürsoy et al. 2017, Perez-Rathke et al. 2020) to generate large ensembles of chromatin fibers within the confines of the cell nucleus without any Hi-C information. These chromatin fibers are modeled as self-avoiding polymer chains consisting of beads, each representing a 5kb genomic region, at the same resolution of Hi-C measurements (Fig1a). These ensembles serve as our background null model. By analyzing measured population Hi-C data and the random null model, we identify a set of statistically significant non-random chromatin interactions that are unlikely to be due to random collisions (Fig1b). Subsequently, we examined these non-random interactions containing genetic variants of eQTLs and extracted relevant information from the reference genome. Further, we use these non-random interactions as input constraints to reconstruct ensembles of 10^5^ single-cell chromatin configurations for selected genomic regions (Fig1c). We then carry out structural analysis on the ensembles of modeled single-cell chromatin configurations to characterize 3D structural properties of the eQTL loci, including conformation compactness, the structure of spatial clusters, and their subpopulation distributions. This provides us with a quantitative assessment of the 3D structural characteristics of the genome and can help us to understand the mechanisms regulating gene expression.

**Figure. 1.**
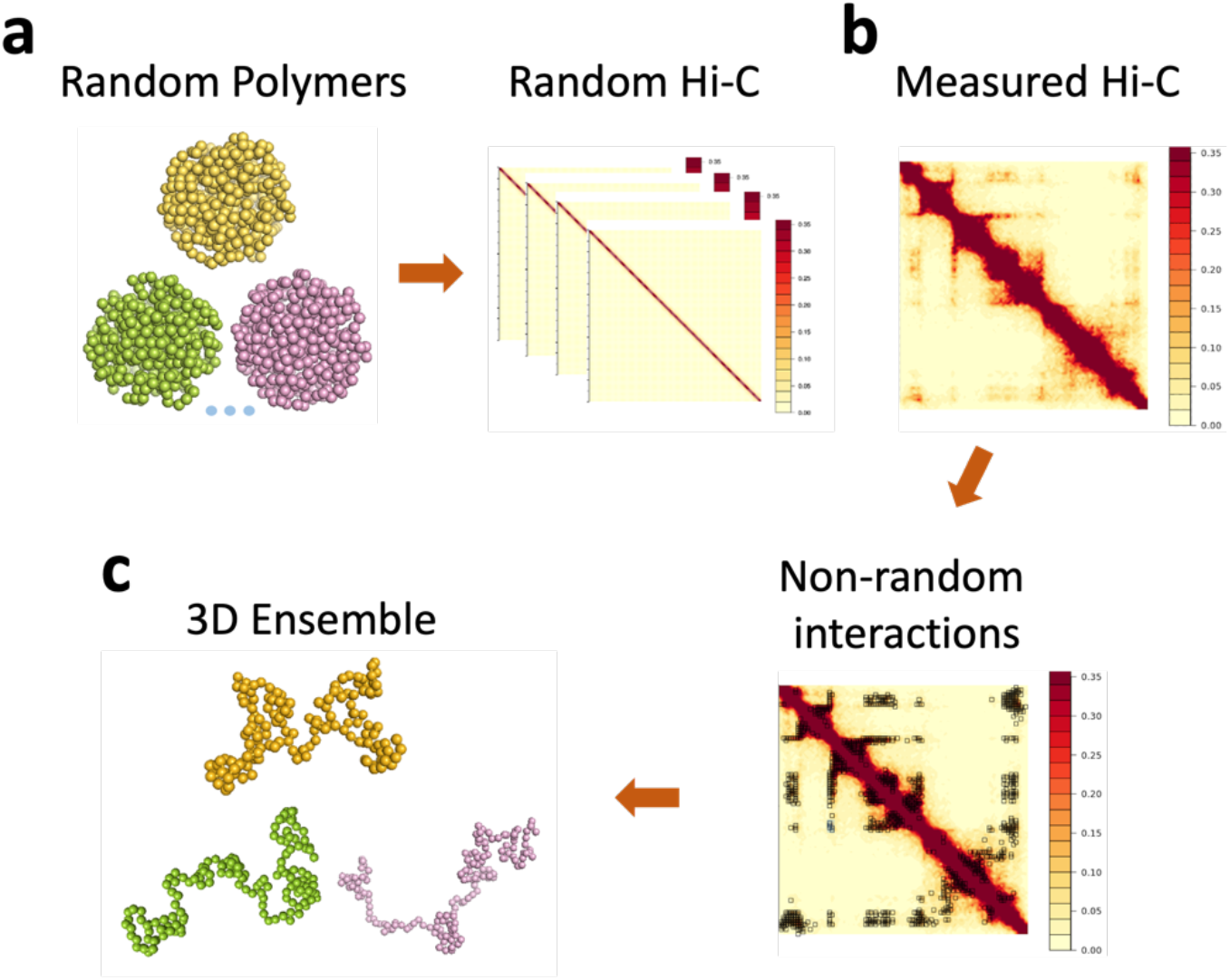
The pipeline of our computational methods. **a.** A large ensemble of random polymer chains is generated as our background null model. **b.** The statistical significance of each Hi-C interaction is computed by bootstrapping the polymer ensemble for identifying non-random interactions from measured Hi-C data. **c.** A large ensemble of single-cell 3D chromatin conformation is reconstructed using the small set of non-random interactions as constrain.

### 2.3 Null model

Our null model was built without any Hi-C data. It comprised 10^6^ random, self-avoiding 3D polymer chains of chromatin fiber within the confines of the cell nucleus. We utilized the fractal Monte Carlo method (Perez-Rathke et al. 2020), which extends the C-SAC technique (Gürsoy et al. 2014, Gürsoy et al. 2017), to construct this model. To address the challenges of sampling in compact space, this process is refined through a recursive resampling algorithm at set checkpoint lengths (Perez-Rathke et al. 2020). The goal is to sample all feasible self-avoiding chromatin chains equally within a defined volume confinement, which is strictly maintained through careful weighting during chain growth (Rosenbluth and Rosenbluth 1955, Grassberger 1997, Liang, Zhang and Chen 2002, Zhang et al. 2003, Liu 2004). The null model consists of 5 × 10^6^ chromatin chains, each 4 Mb in length and is made up of 800 beads at 5 kb resolution. The volume confinement is proportional to the nuclear volume of cell types.

### 2.4 Non-random interactions

By comparing the Hi-C measurements to our null model, we were able to distinguish between the non-random interactions that may contribute to the 3D chromatin structures and the by-stander interactions that most likely are due to random polymer collision. To estimate the random contact probabilities of pairs of loci, we determine the frequency of 3D configurations where the spatial distance between the corresponding bead pair is less than 80 nm (Giorgetti et al. 2014, Gürsoy et al. 2017). The statistical significance of each Hi-C interaction is determined through bootstrapping the random ensemble (Kleiner et al. 2014). Hi-C interactions with a BH-FDR adjusted *p*-value below 0.05 are considered as non-random interactions (Benjamini and Hochberg 1995).

### 2.5 3D single-cell chromatin conformations

By taking these non-random interactions as physical constraints, we generate a large ensemble (5 × 10^4^) of folded 3D chromatin conformations. All conformations are constructed by using our previously described chromatin folding algorithms, which are based on a novel approach of sequential Bayesian inference (Perez-Rathke et al. 2020, Sun et al. 2021).

## 3. RESULTS

### 3.1 A small fraction of Hi-C contacts are non-random interactions

It is well known that many of the Hi-C measurements may be by-stander contacts resulting from random collisions of chromatin fibers within the limited space of the nucleus(Belmont 2014, Gürsoy et al. 2017). Our null model of random polymer ensumbles can help to eliminate those spurious by-stander contacts and identify important Hi-C contacts that contribute to the formation of 3D chromatin structure.

We take Hi-C data of three loci from lymphoblastoid cells (GM12878), human primary mammary epithelial cells (HMEC) and lung fibroblast cells (IMR-90) from 4DN. Each loci containing eQTLs variants associated with ≥ 1 gene for at least one of these tissues. These loci are chr2: 230,805,000-231,690,000 (Locus I), chr16: 85,845,000-86,580,000 (Locus II) and chr18:10,530,000-11,160,000 (Locus III).

Our results show that just a fraction of Hi-C interactions can be considered as non-random. Specifically, 9.8%, 8.9%, and 13.1% of the total contacts for GM12878 cells are identified as non-random interactions for these three loci (Figure 2). 10.1%, 12.9%, and 14.9% of the total contacts for IMR90 cells are identified as non-random interactions and 13.4%, 22.4%, and 31.1% of the total contacts for HMEC cells are identified as non-random interactions (see Supplementary Figure 1). They all meet the specified criteria that adjust *p*-value is less than 0.05. The high proportion of non-random interactions in HMEC cells is due to the low read count of the HMEC Hi-C data. Although they constitute of only a small fraction of Hi-C interactions, the key structural patterns in the heatmap are well-presented by the non-random interactions in all three loci (Figure 2).

**Figure. 2.**
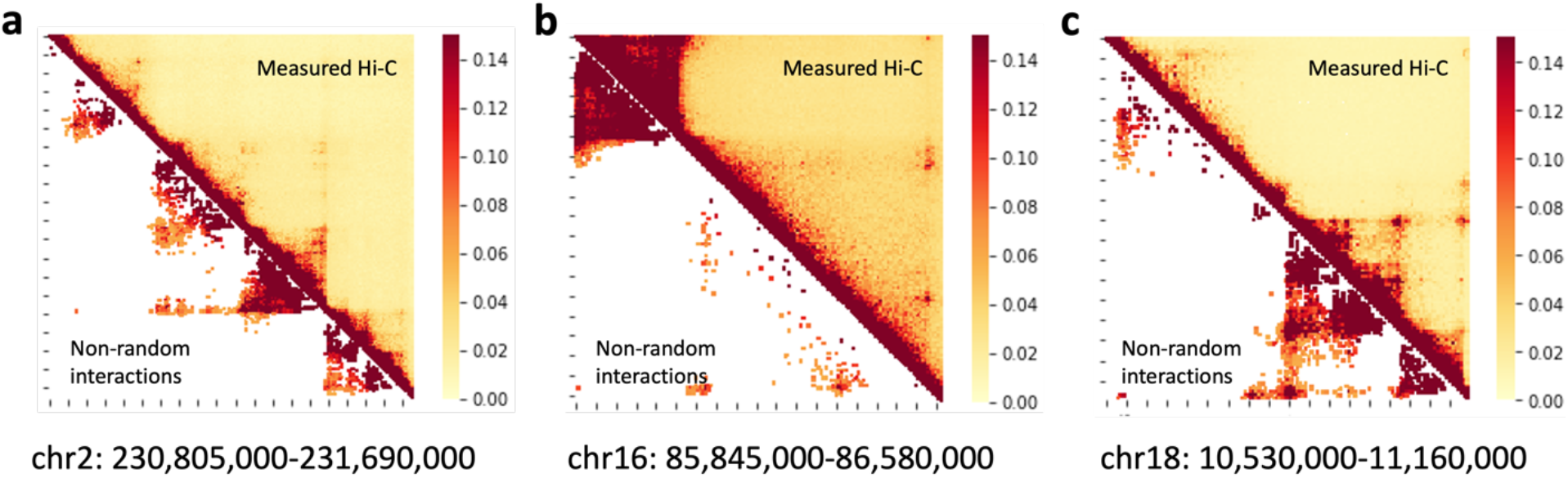
Small fractions of Hi-C contacts are non-random interactions in GM12878 cells. The non-random interactions of loci a. chr2: 230,805,000-231,690,000, b. chr16: 85,845,000-86,580,000 and c. chr18: 10,530,000-11,160,000. The percentage of non-random interaction is 9.8%, 8.9% and 13.1%, respectively.

### 3.2 Non-random interactions can reconstruct ensembles of single-cell chromatin conformation

To evaluate the importance of these non-random interactions we uncovered, we use these non-random interactions as physical constraints to generate 3D chromatin conformations. We used previously developed efficient chromatin folding algorithms (Sun et al. 2021).

For Locus I, there are 10%-13% of the total contact pairs in GM12878, IMR90, and HMEC (Figure 3a). Using these non-random interactions, we reconstruct large ensembles of 5 × 10^5^ single-cell chromatin conformations for each cell line at 5kb resolution. To compare our predicted ensemble of single-cell chromatin conformation with the experiment-measured Hi-C data, we calculate the interaction probabilities of simulated chromatin conformations. Taking the basic premise that DNA segments near each other are potential candidates for Hi-C ligations, we determined the interaction probabilities based on the frequency of 3D structures where the 3D distances between certain pairs of loci are less than 80 nanometers (Giorgetti et al. 2014, Gürsoy et al. 2017). As shown in Figure 3b, the simulated Hi-C map, obtained by aggregating 10,000 single-cell conformations, exhibits strong similarities with the measured Hi-C maps for GM12878, IMR90, and HMEC for Locus I with Pearson correlation coefficients range from 0.94 to 0.97. Similar results are observed for the other two loci, where Pearson correlation coefficients are 0.95, 0.9, and 0.96 for Locus II, and 0.96, 0.87, and 0.95 for Locus III. (see Supplementary Figure 2-3)

**Figure 3.**
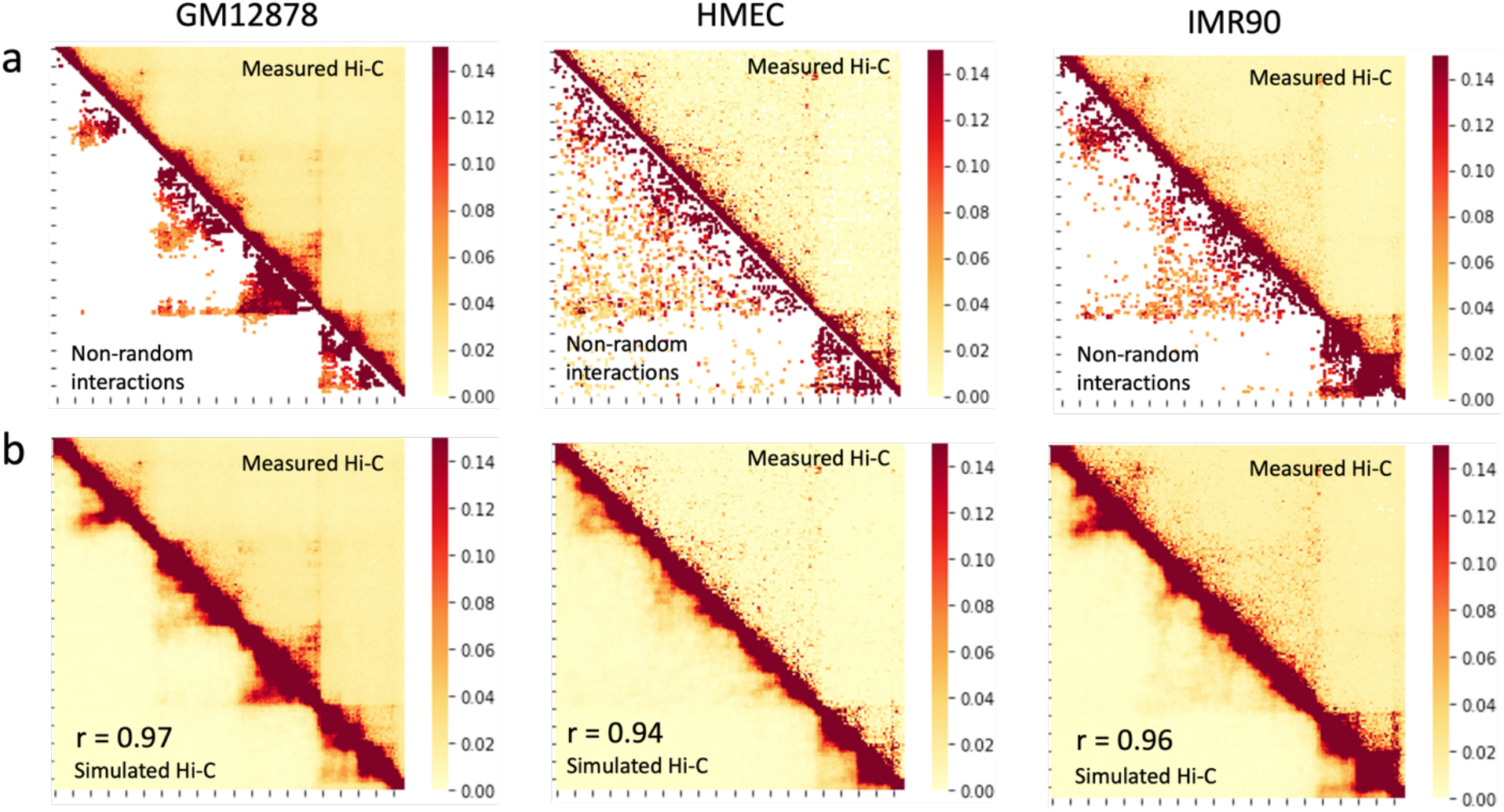
Precise reconstruction of 3D chromatin conformations with high resolution **a**. Non-random interactions identified from our model for GM12878, HMEC, and IMR90 cell lines in Locus I. **b**. Simulated Hi-C map compared to the measured Hi-C map for GM12878, HMEC, and IMR90 cell lines within Locus I. The corresponding Pearson correlation coefficients (R) are 0.97, 0.96, and 0.94, respectively.

These findings demonstrate that by utilizing just a fraction of the Hi-C interactions identified as non-random interactions, we can accurately reconstruct experimental Hi-C contact heatmaps for different regions of chromosomes at high resolution with accuracy. Our results show that the non-random interactions we uncovered are sufficient to guide chromatin folding of these loci in all three cell types.

### 3.3 Uncovering physical contacts between eGene and eQTLs and their strong influence on gene expression

Previous studies showed that eQTLs tend to be located near their target genes in space and loci containing eQTLs show a higher frequency of contact interactions (Duggal, Wang and Kingsford 2014). Additionally, eQTLs enriched with cis-regulatory elements are commonly situated in close spatial proximity to the promoters of the genes they target (Aguet et al. 2017, Duggal, Wang and Kingsford 2014). With the 3D single-cell chromatin conformations now at hand, we examined the 3D structures of our selected eQTL containing loci.

We first analyzed modeled single-cell conformations to determine the occurrence where two 5kb genomic regions represented by polymer beads, one containing eGene and another containing eQTLs, are in physical interaction using the criteria that the median distance between them is less than 80 nm (Giorgetti et al. 2014, Gürsoy et al. 2014). Next, we examine all eGene-eQTLs pairs derived from the GTEx database to determine whether they are in physical contacts among all three loci for EBV-transformed lymphocytes (GM12878), breast–mammary tissue (HMEC), and lung cells (IMR90). eGene-eQTLs pairs are then categorized as either in physical contact or are free of such contacts.

Among the three loci, we found 153 physical eGene-eQTLs contacts from 798 eGene-eQTLs pairs in EBV-transformed lymphocytes, 365 physical eGene-eQTLs contacts from 2,024 eGene-eQTLs pairs in Brest-mammary tissue, and 535 physical eGene-eQTLs contacts from 2,678 eGene-eQTLs pairs in lung tissue cells. Our method can reveal pairs of eGenes and eQTLs in physical contact. (For more details on these physically contacted eGene-eQTL pairs, see Supplementary Data.)

Intriguingly, our results show that physically contacting eQTLs exhibit a stronger influence on eGene than those without physical contact, as they have larger normalized effect size (NES) values (Lonsdale et al. 2013). Among the three tissues, physically contacting eGene-eQTLs pairs have larger absolute values of NES than non-physically contacted pairs (Figure 4). Here, the NES value of an eQTLs is defined as the slope of the linear regression and is taken as a measure of the effect of the alternative allele (ALT) relative to the reference allele (REF) in the human genome reference following (Lonsdale et al. 2013). A higher normalized effect size suggests that the genetic variant has a stronger influence on gene expression (Lonsdale et al. 2013, Zeng et al. 2022). Among all three tissues, the NES absolute values difference between physical contact pairs and non-physical contact pairs is significant, which the *p*-value is 1.0×10^-3^, 3.05×10^-11^ and 2.2×10^-2^, respectively.

**Figure 4.**
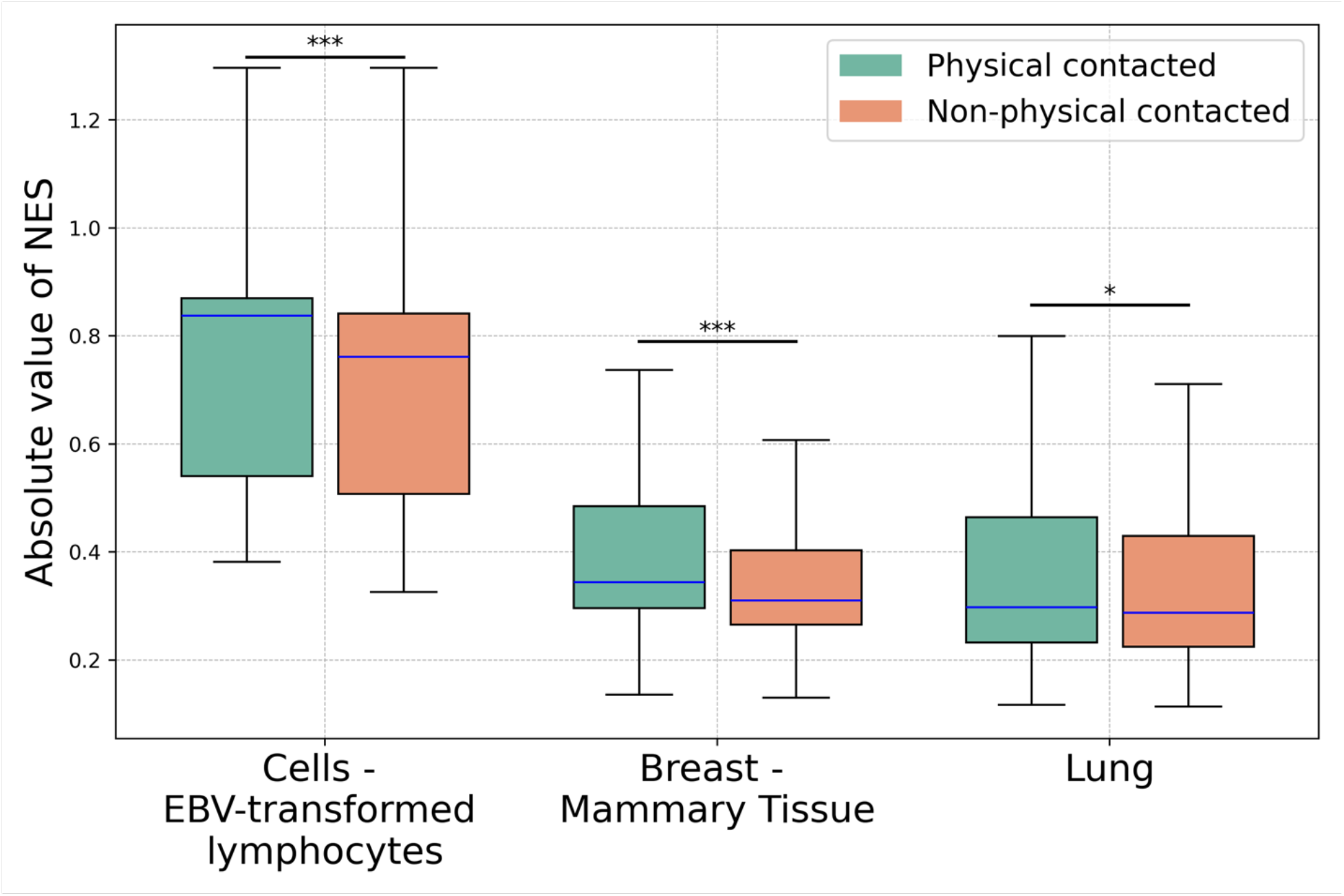
Display of the absolute values of NES for eGene-eQTL pairings across all three examined loci for EBV-transformed lymphocytes (GM12878), breast - mammary tissue (HMEC) and lung (IMR90) cells, respectively. A larger absolute NES value indicates a more substantial genotype impact on gene expression. Significance levels are indicated as follows: ***p < 0.001, **p < 0.01, *p < 0.05.

### 3.4 Discovery of many-body interactions at eGene and eQTLs loci

Previous studies found that many-body chromatin interactions such as those in condensates likely play important roles in gene regulations (Hnisz et al. 2013, Gürsoy et al. 2017, Sabari et al. 2018, Quinodoz et al. 2018, Perez-Rathke et al. 2020). However, the presence of many-body interactions within individual cells are masked in the bulk Hi-C measurements, as they capture average pairwise genomic interactions across populations. In order to determine whether many-body interactions occur in eGenes and eQTLs loci, we investigate Locus II (chr16: 85,845,000-86,580,000) across GM12878, HMEC and IMR90 cells, which contains eGenes RP (RP11-463O9.9) and MT (MTHFSD), both are associated with the eQTL of chr16_86531581_A_T_b38 (Figure 5a, b). We examined the spatial structure of genetic interactions among these eGenes and eQTL at this locus.

**Figure 5.**
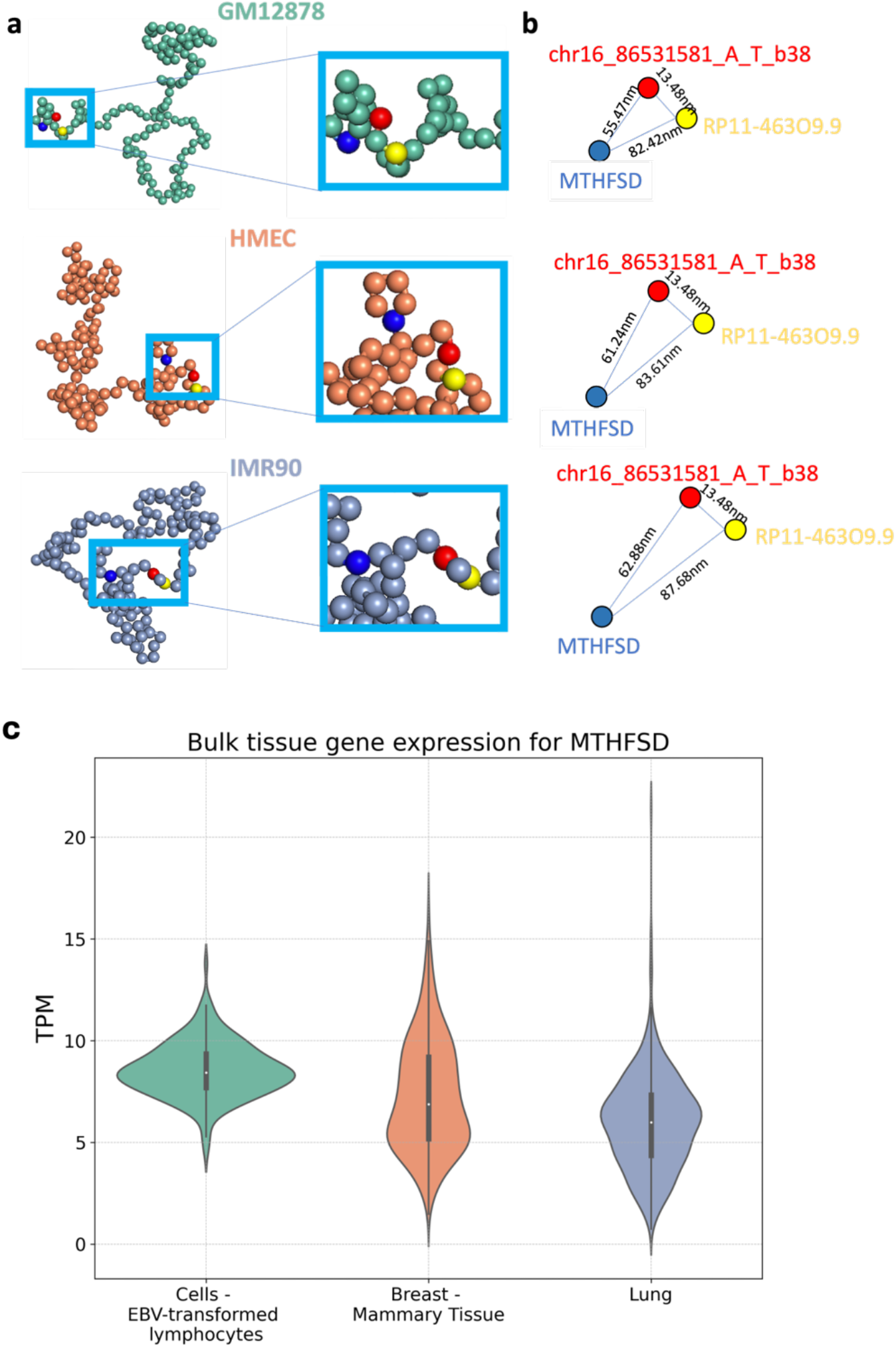
The physical interactome of many-body genetic interactions between eGene and eQTL at Locus II. **a**. Representative spatial structures of the locus for lymphocytes (GM12878), breast-mammary tissue (HMEC) and lung fibroblasts (IMR90), with eGenes MTHFSD (blue), RP11-463O9.9 (yellow) and eQTLs chr16_86531581_A_T_b38 (red) depicted. Note the spacial distances among eGenes and eQTL are different. **b**. The spatial distances between the eGenes MTHFSD (blue), RP11-463O9.9 (yellow), and the eQTLs (red) of the many-body interaction unit in the representative structures shown in (**a**). The spacial distances among the many-bodies are the closest in GM12878, followed by HMEC and then IMR90 **c.** Bulk tissue gene expression level of eGene MTHFSD among the three tissues of lymphocytes (GM12878), breast-mammary tissue (HMEC), and lung (IMR90). The expression level of MTHFSD is the highest in the transformed lymphocytes cells, followed by breast-mammary cells and then lung cells.

Analysis of the 3D chromatin structure shows that three-body interactions do exist at this locus. Here, we define three-body interactions as a complex of 5kb chromatin regions, where the median Euclidean distances between all pairs of regions in the complex of the ensemble of chromatin conformations are shorter or approximately equal to a cross-linking threshold of 80 ± 5 nm (Giorgetti et al. 2014, Gürsoy et al. 2017) (Figure 5b). The interactomes among eGenes RP, MT and the eQTL are three-body in both GM12878 and HMEC. However, the degree of influence of the eQTL variant on gene expression is different among these three tissues. This is indicated by the different values of NES for the same eQTL. The absolute value of NES of this eQTL on eGene MT in lymphocytes (GM12878) is the largest (1.9). The NES values are only 0.64 for breast-mammary tissue (HMEC) and 0.53 for lung tissue (IMR90), respectively. Furthermore, the NES of the same eQTL with eGene RP is also the largest (1.6) in lymphocytes (GM12878). The NES value is 1.4 for breast-mammary tissue (HMEC) and 1.0 for Lung tissue (IMR90), respectively.

To understand the reason why the impact of the variant on gene expression as measured by NES value varies across these three tissues, we ask whether it can be explained by their spacial distance relationship. For this, we randomly selected 5,000 simulated single-cell conformations from the ensembles for GM12878, HMEC, and IMR90 cells and calculated the Euclidean distance between eQTL and the eGenes. From the distributions of the Euclidean distance between eGene and eQTLs, we found that in GM12878 the median spatial distance between the promoter of the eGene MT and the eQTL is 55.74 nm. The distance between the two promoters of the eGenes MT and RP is 82.42 nm. These are closer than the distances in HMEC (61.24 nm, 83.61 nm) and IMR90 (62.88 nm, 87.68 nm). For eGene RP, the spatial distance between its promoter and the eQTL is similar among all three cell types; this is consistent with their shared close genomic distances between the eQTL and the eGene RP (<10kb). The shorter 3D physical distances in GM12878 (MT to the eQTL and MT to RP) among the eGenes MT, RP and the eQTL is closer than the corresponding 3D physical distance in HMEC and in IMR90, leading to stronger effects of eQTL on gene expression in GM12878. These results demonstrate that closer spatial proximity between eQTL and the target gene promoters can result in stronger eQTL effects on gene expression, as revealed by the larger absolute value of NES.

The measured expression level of eGene MT in these tissues provide additional direct support to our findings (Figure 5c). When eQTL has closer spatial distance to MT in GM12878 cells (55.73 nm vs 61.24nm in HMEC and 62.88 nm in IMR90), it has stronger effects to the gene expression: The median TPM of gene MT in lymphocytes is the largest at 8.55, followed by 7.27 in breast-mammary tissue and 5.97 in lung tissue. Overall, these results show that there exist three-body interactions among eGenes RP, MT and eQTL at Locus II, and the spatial pattern of interactions among the eQTLs and eGenes play important roles in gene regulation and genome functions.

### 3.5 3D chromatin structure difference and heterogeneities at eGene and eQTLs loci among tissues

Quantitatively assessing the chromatin heterogeneities of cells within a tissue is challenging, as chromatin in individual cells may take different configurations, and the overall ensemble may be very heterogeneous. Because of this, initial condition independent thorough sampling is crititcal. With a large and diverse ensemble of single-cell conformations generated, we can group them into clusters by their 3D topology to quantify their heterogeneity (Perez-Rathke et al. 2020, Sun et al. 2021). We use the K-means clustering method to cluster the 3D chromatin configurations into major subpopulations of chromatins with similar topologies. For Locus I, we used 10,000 simulated single-cell chromatin conformations for each of GM12878, IMR90, and HMEC cells, totaling 30,000 conformations, and were able to group them into four clusters (Figure 6). There are notable distinctions in both the contacts heatmap and compactness among the four clusters. Importantly, the distribution of subpopulations varies significantly across different cell lines, with HMEC and IMR90 predominantly featuring subclusters 1 and 3, respectively, while GM12878 enriched with subclusters 2 and 4 cells. That is, GM12878 cells are more heterogeneous compared to HMEC and IMR90 cells. Our simulated chromatin conformations uncovered significant structural differences among different tissues. Similar results are also observed for Locus II and Locus III (see Supplementary Figure 4-5). These findings highlight substantial differences in the conformation of different cell lines in the same locus and reveal strong tissue-specific 3D chromatin structures, which are not evident in the population Hi-C heatmap directly (Figure 6).

**Figure. 6.**
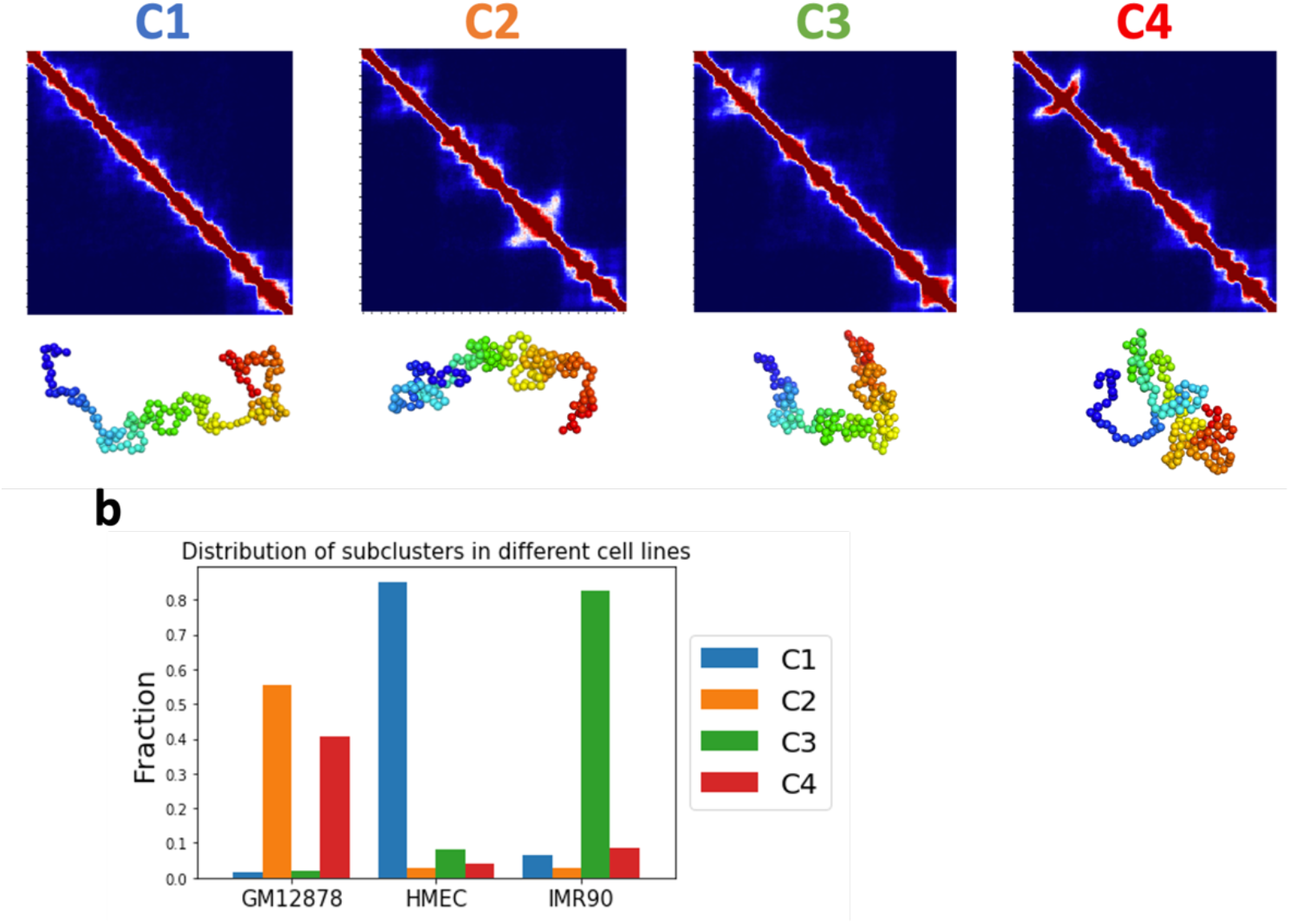
Uncovering the tissue-specific subpopulations of 3D chromatin structures. **a** 3D chromatin conformation of GM12878, IMR90 and HMEC at Locus I (chr2: 230,805,000-231,690,000). The resulting aggregated contact heatmaps of the four clusters are arranged in order of their compactness. Representative conformations of four clusters are shown below the heatmap. b Proportions of the four subpopulations in each cell types.

## 4 CONCLUSION AND DISCUSSION

In this study, we have developed a set of tools to study the relationship between eQTLs and 3D genome organization, through modeling of spatial and physical chromatin interactions at the single-cell level, enabling a general understanding of how eQTLs affect gene expression. Our method identified a small set of non-random interactions from the measured Hi-C data that effectively captured the key patterns from the population Hi-C heatmap. Using these non-random interactions, we reconstruct the 3D single-cell chromatin conformations with accuracy. These simulated chromatin conformations provide detailed structural insights that enable the examination of the 3D spatial pattern of chromatin at loci containing eQTLs and eGenes.

Furthermore, our work enables new discoveries as we can now directly investigate the 3D spatial relationship between eGenes and eQTLs across multiple tissues. Results obtained from the analysis of 3D single-cell chromatin conformations of loci across different cell types showed that the physically interacted eQTLs have stronger effects on gene expression. The specific spatial arrangements of eQTLs in cell subpopulations allow a quantitative approach to understanding the regulatory mechanisms controlling gene expression. In addition, our tools enable novel and compelling biological questions to be formulated, such as possible roles of specific promoters, enhancers, super-enhancers, genes, and other elements relating to specific eQTLs in multi-way interactions.

While we have shown a promising approach to investigate genome structure and function relationship, there are limitation. Firstly, the availability of high-quality Hi-C data is limited. With more high-quality data from the 4DN becoming available recently, this will become less of an issue. Secondly, we analyzed only three loci among three cell lines. It is desirable to expand the scale to analyze additional eQTLs and eGene-associated loci among many more tissues and cell lines. Furthermore, the advancement of single-cell sequencing technologies, such as scRNA-seq (Tang et al. 2009), scATAC-seq (Buenrostro et al. 2015), and scHi-C (Ramani et al. 2017) provided additional data at single cell level. Integrating these data with 3D chromatin modeling should lead to better understanding of genome structure-function relationship at the single-cell level.

## Acknowledgment

This work is supported by NIH grants 1R03OD032628-01, 1R03OD036492-01 and 2R35GM127084-06. An award for computer time was provided by the U.S. Department of Energy’s (DOE) Innovative and Novel Computational Impact on Theory and Experiment (INCITE) Program. This research used resources from the Argonne Leadership Computing Facility, a U.S. DOE Office of Science user facility at Argonne National Laboratory, which is supported by the Office of Science of the U.S. DOE under Contract No. DE-AC02-06CH11357.

## Data availability

Non-random interactions, single-cell conformations and physically contacted eGene-eQTLs pairs are available on UIC Indigo (https://indigo.uic.edu/projects/Constructing_High-Resolution_Ensemble_Models_of_3D_Single-Cell_Chromatin_Conformations_of_eQTL_Loci_from_Integrated_Analysis_of_4DN-GTEx_Data_towards_Structural_Basis_of_Differential_Gene_Expression/190164)

