## Supplemental figures for "Structural Basis of Differential Gene Expression at eQTLs Loci from High-Resolution Ensemble Models of 3D Single-Cell Chromatin Conformations"

Supplementary Figures

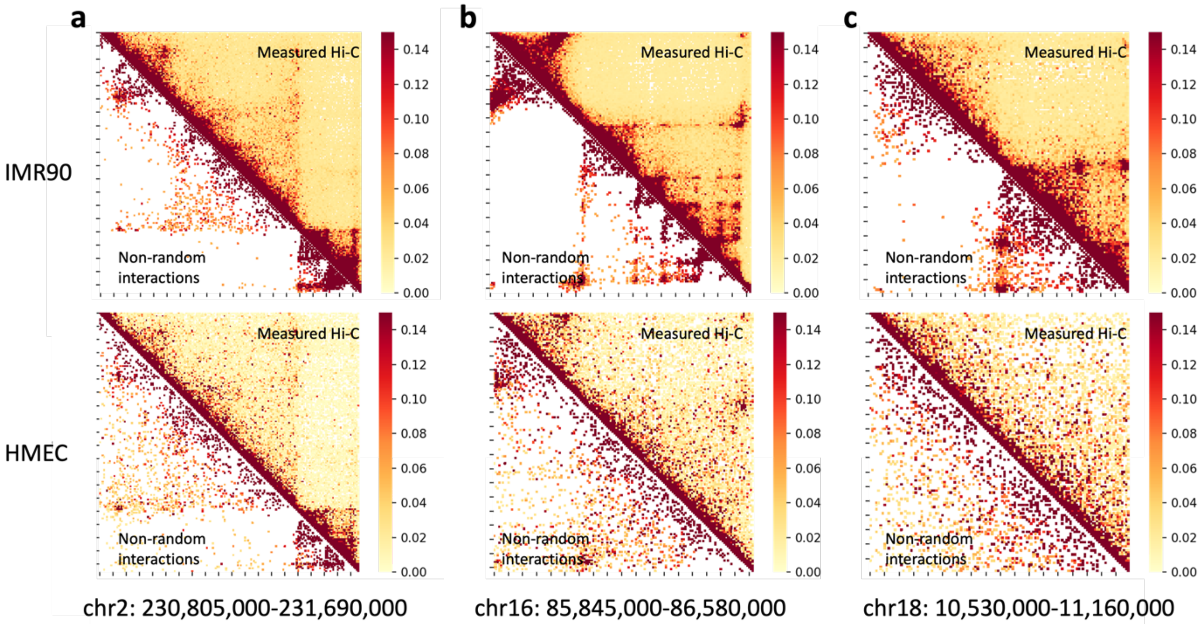


**Supplementary Figure 1.** Small fraction of Hi-C contacts are non-random interactions. The non-random interactions of loci **a.** chr2: 230,805,000-231,690,000, **b.** chr16: 85,845,000-86,580,000 and **c.** chr18: 10,530,000-11,160,000 in IMR90 and HMEC. The percentage of non-random interaction is 10.2%, 12.9% and 14.9% in IMR90 and 13.4%, 22.4%, and 31.1% in HMEC separately.

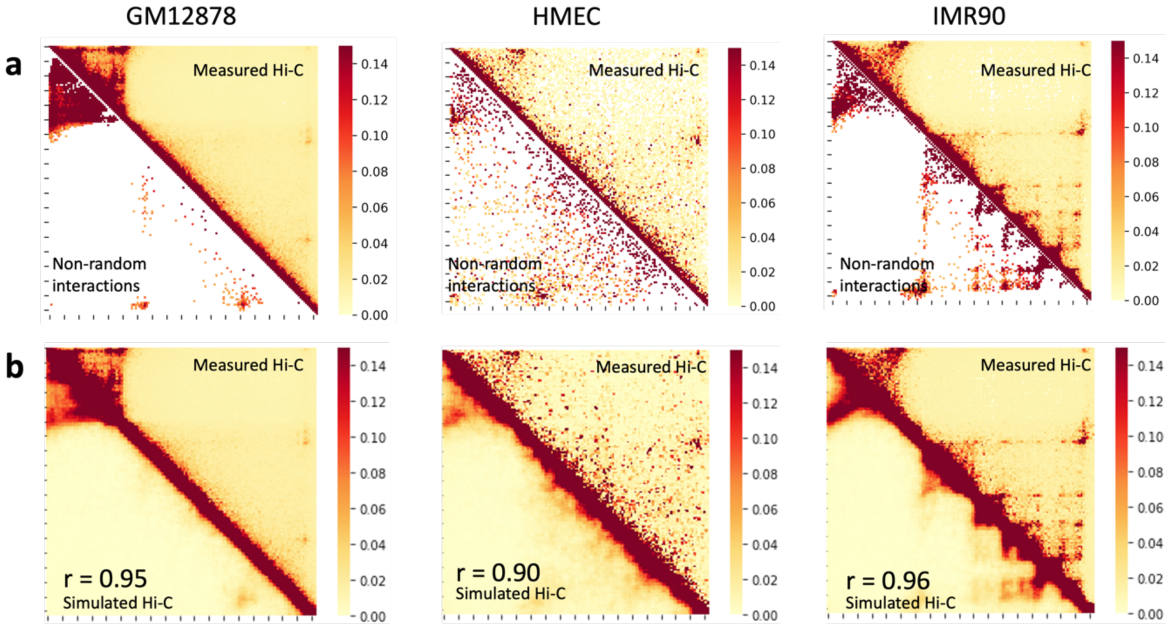


**Supplementary Figure 2.** Precise reconstruction of 3D chromatin conformations with high resolution **a**. Non-random interactions identified from our model for GM12878, HMEC, and IMR90 cell lines within the genomic region chr16: 85,845,000-86,580,000. **b**. Simulated Hi-C map compared to the measured Hi-C map for GM12878, HMEC, and IMR90 cell lines within the genomic region chr16: 85,845,000-86,580,000. The corresponding Pearson correlation coefficients (r) are 0.95, 0.90, and 0.96, respectively.

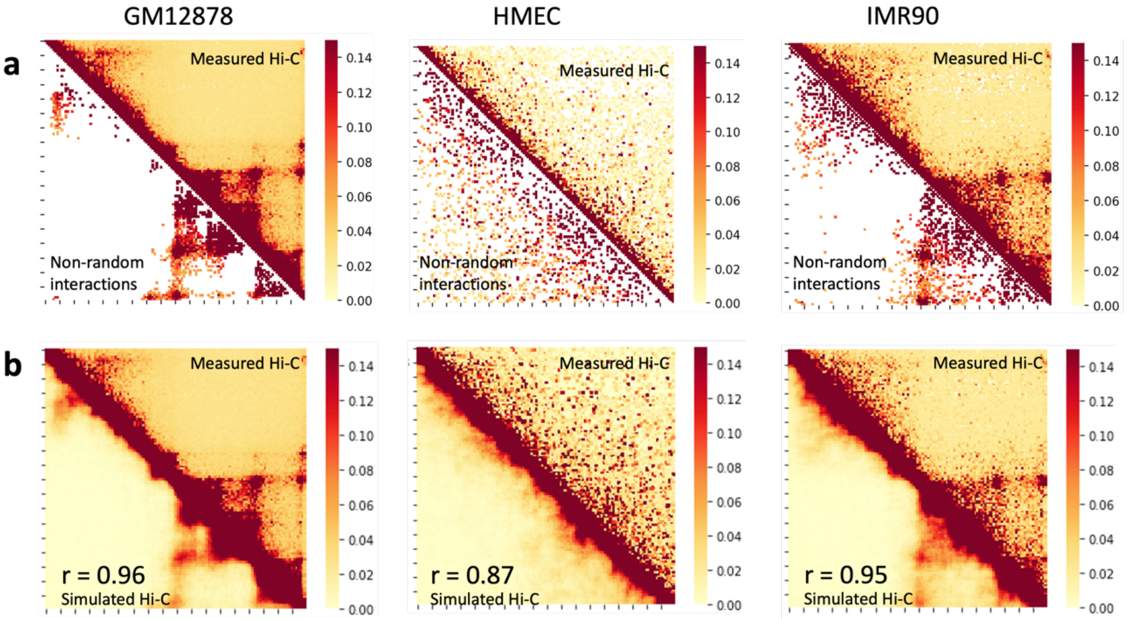


**Supplementary Figure 3.** Precise reconstruction of 3D chromatin conformations with high resolution **a**. Non-random interactions identified from our model for GM12878, HMEC, and IMR90 cell lines within the genomic region chr18: 10,530,000-11,160,000. **b**. Simulated Hi-C map compared to the measured Hi-C map for GM12878, HMEC, and IMR90 cell lines within the genomic region chr18: 10,530,000-11,160,000. The corresponding Pearson correlation coefficients (r) are 0.96, 0.87, and 0.95, respectively.

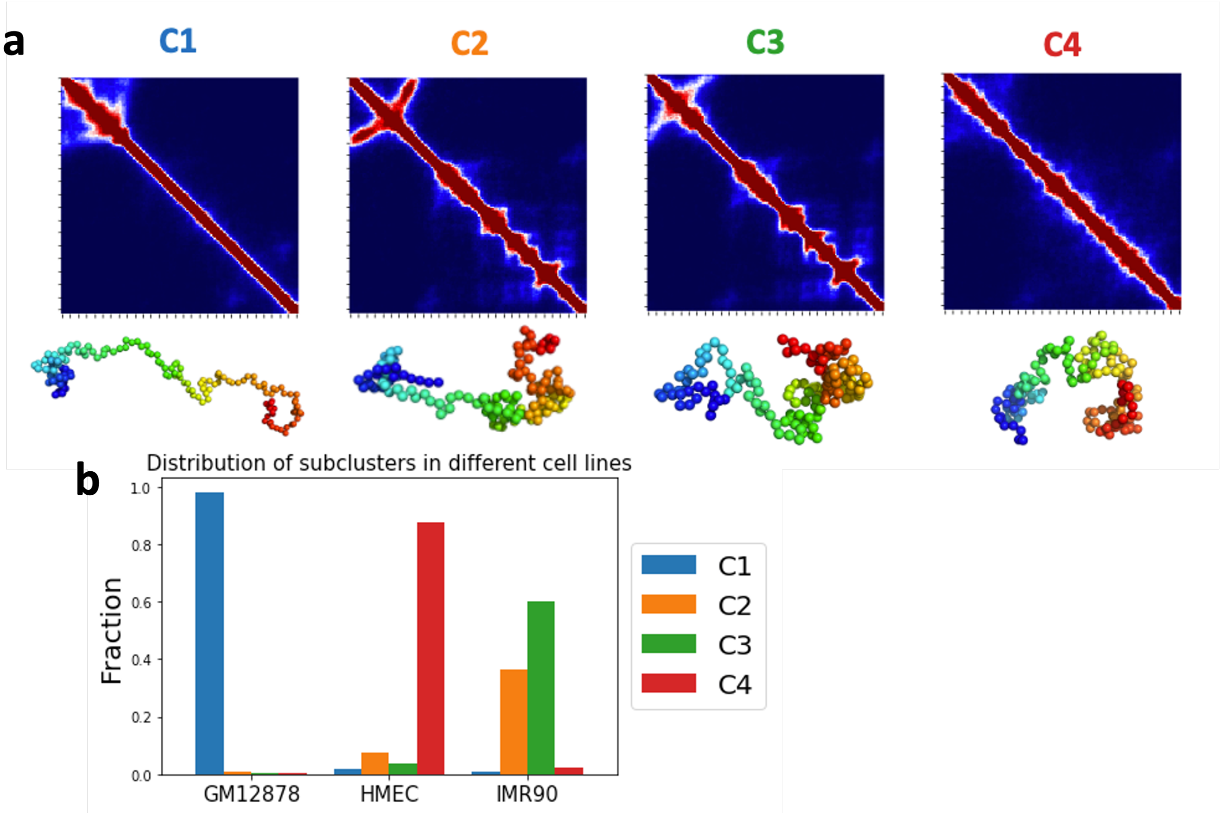


**Supplementary Figure 4.** Chromatin subpopulation uncover the tissue-specific 3D chromatin structure. **a** 3D chromatin conformations of GM12878, IMR90 and HMEC in genomic locus chr16: 85,845,000-86,580,000. The resulting aggregated contact heatmaps of the 4 clusters are arranged in order of their compaction. Representative conformations of 4 clusters shown below the heatmap. **b** Proportions of the 4 subpopulation in each cell types.

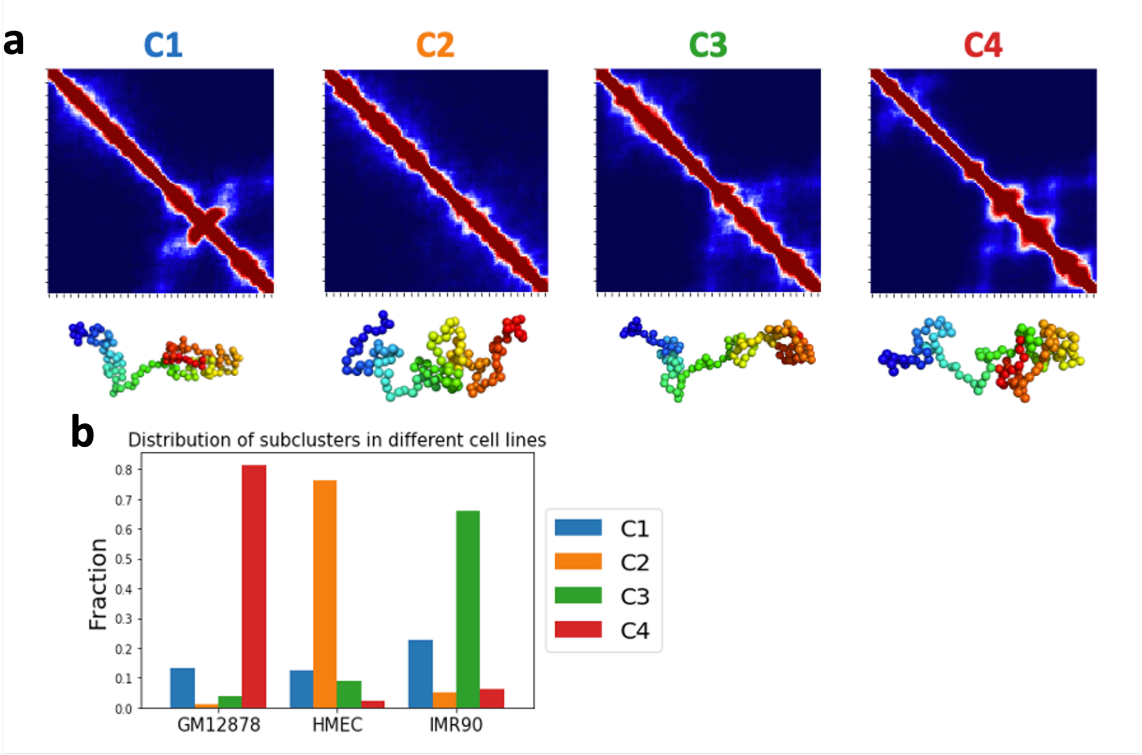


**Supplementary Figure 5.** Chromatin subpopulation uncover the tissue-specific 3D chromatin structure. **a** 3D chromatin conformations of GM12878, IMR90 and HMEC in genomic locus chr18: 10,530,000-11,160,000. The resulting aggregated contact heatmaps of the 4 clusters. Representative conformations of 4 clusters shown below the heatmap. **b** Proportions of the 4 subpopulation in each cell types.
